# Future heatwave conditions inhibit CO_2_-induced stomatal closure in wheat

**DOI:** 10.1101/2025.01.17.633535

**Authors:** Robert S. Caine, Muhammad S. Khan, Yixiang Shan, Colin P. Osborne, Holly L. Croft

## Abstract

Rising atmospheric CO_2_ concentrations are driving ongoing climatic changes, leading to agricultural crops increasingly experiencing extreme weather events^1^. Stomata serve as gatekeepers on plant leaves, regulating both CO_2_ capture for photosynthesis and the concomitant release of water. At higher CO_2_ concentrations or higher vapour pressure deficit (VPD), stomatal pores narrow, reducing stomatal conductance to water vapour (*g_sw_*) and transpiration (*E*)^2–6^. Increasing temperatures and/or nitrogen fertilisation promote an opposite stomatal response, enhancing *g_sw_* and *E*^7,8^. With atmospheric CO_2_ concentration, temperature and VPD predicted to rise throughout this century^1^, it is unclear how crops will modify stomatal gaseous exchanges, particularly under differing N-fertilisation regimes. Here, we show in wheat (*Triticum aestivum*), that elevated CO_2_ does not reduce *g_sw_* or *E* during heatwaves when VPD is high, instead plant water usage increases. High-VPD heatwave events also impact stomatal responsiveness to N-fertiliser application, prompting significantly higher gas exchange contributions from abaxial leaf surfaces, irrespective of CO_2_ growth conditions. Dynamic stomatal responsiveness to light and high CO_2_ are also attenuated during heatwaves in a CO_2_-independent manner. Taken together, our data suggests that future wheat crops will use significantly more water during heatwaves than might be expected, which has substantial implications for future global food security.

## Main

The global atmospheric CO_2_ concentration is currently 425 ppm^9^, having risen from ∼280 ppm during pre-industrial times^10,11^. Forecasts suggest that CO_2_ increases will continue throughout much of the century, with Shared Socioeconomic Pathways (SSP) predictions ranging from 393 ppm (SSP1-1.9) to 1195 ppm (SSP5-8.5) by 2100^1,12^. Without a unified global stance on climate action, it is unclear which SSP is most likely to unfold by the end of the century, and if fossil fuel usage rates continue as they are now, CO_2_ levels could reach 620-860 ppm^12,13^. Many plants benefit from increased CO_2_ fertilisation due to enhanced photosynthesis (*A*) and reduced *g_sw_* promoting greater intrinsic water-use efficiency (iWUE) and drought tolerance^14,15^. However, with increasing atmospheric CO_2_ expected to co-occur with rising temperatures and higher VPD, it is unclear what the net impacts on plant gaseous exchanges will be, particularly because *E* is fundamental for evaporative cooling^16^ and for obtaining nutrients required to support higher rates of *A*^17,18^.

Wheat provides 20% of daily calories worldwide and is the largest crop globally by cultivated land area^19–21^. Yields of wheat have increased substantially over the last century, thanks in large part to increased application of inorganic nitrogen (N) fertiliser application^22,23^, but this has been at the expense of increased water usage^24^; with highly fertilised wheat crops considerably more susceptible to drought^25,26^. Extreme heatwave events also raise wheat water requirements^27^, often resulting in drought and reduced fertility as temperatures become increasingly severe^28,29^. Currently, drought and heat-related losses account for a 12.4% reduction in global wheat yields^30,31^, but this yield gap is expected to grow as climate conditions become increasingly extreme^32^. This is particularly concerning as water demand will outstrip supply by ∼40% by 2030^1^. With demand for food continuing to grow throughout much of this century^33^, evidence suggests that it will be increasingly difficult to meet demands. It remains unclear if rising atmospheric CO_2_ concentration can offset the expected global water deficit of agricultural crops to mitigate yield losses, particularly given the predicted increases in extreme weather events such as heatwaves.

Here, we investigate the interactive effects of rising atmospheric CO_2_ concentration and higher VPD on wheat water usage, resilience and overall productivity. Conducting experiments across a gradient of N-fertiliser treatments, we ask the following questions: (1) Will higher atmospheric CO_2_ concentration lead to reduced water usage during high-VPD heatwaves? (2) Do stomata respond differently to light, CO_2_ and N-fertiliser application when grown under differing CO_2_ concentrations and VPD? And (3) what are the overall impacts on wheat water usage and productivity, when high CO_2_, high VPD, drought and varying N-fertiliser are considered in combination?

### High CO_2_ promotes cooling at high VPD

Experimental scenarios simulating rising global CO_2_ and VPD concentrations at higher temperatures, across a range of high-N fertiliser application are presented in **Fig. 1**. Plants were grown at 450 ppm (ambient) or 720 ppm (high) CO_2_ concentration (2 experiments per CO_2_ treatment), with a 16-day heatwave treatment (27 °C, 60% RH → 37 °C, 50% RH) applied to 1x each CO_2_ experiment, starting when flag leaves first emerged. The heatwave treatment increased VPD from 1.43 kPa (ambient) to 3.14 kPa (high). To explore if high CO_2_ mitigated the effects of high VPD treatment during water-deficit, a drought treatment was applied for 12 days (9 days into heatwave treatments), starting as wheat ears were emerging from canopies. For plants not exposed to heatwave treatments, drought was imposed slightly later due to slower phenological development.

**Fig. 1:**
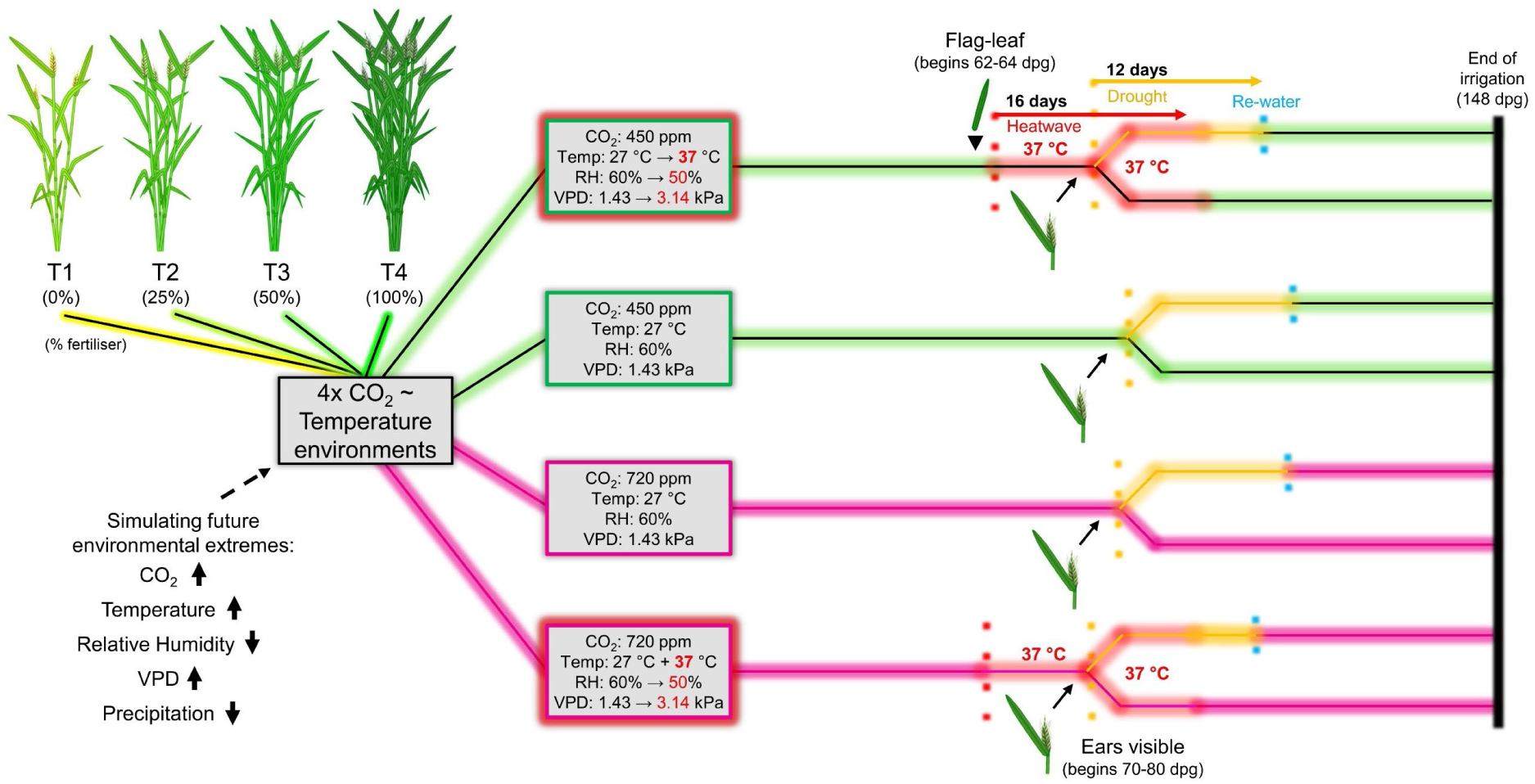
Simulating future environmental extremes to study wheat responses to rising CO_2_ and high vapour pressure deficit (VPD) heatwaves. Wheat was cultivated under either 450 ppm or 720 ppm CO_2_ concentration, with VPD set to 1.43 kPa and then increased to 3.14 kPa for heatwave treatment as flag leaves began appearing. Heatwaves were applied for 16 days, with a 12-day drought period imposed from 9 day into the heatwave treatment, as wheat ears were emerging. A range of high N-fertiliser treatments were included within each growth scenario (T1 = no high N-fertiliser application → T4 = maximum high N-fertiliser application). *n = 32* per fertiliser treatment per scenario, reducing to 16 during drought.

Previous research shows that higher atmospheric CO_2_ concentrations often induce stomatal closure leading to reduced *E* and *g_sw_* rates^2,3^. By measuring leaf *E* and *g_sw_* 5 days into high-VPD heatwaves, we explored wheat water flux responses to elevated CO_2_ and/or high VPD exposure, across a range of high-N fertiliser treatments (**Fig. 2**, Extended Data Fig. 2). During ambient VPD, non-heatwave conditions, high CO_2_ grown wheat displayed lower combined leaf *E* (abaxial and adaxial surfaces combined) than ambient CO_2_ equivalent plants across all N treatments (**Fig. 2a**). Strikingly, high-VPD heatwave exposure resulted in the opposite response in 3 out of 4 N-fertiliser treatments, with high CO_2_ N-fertilised plants having higher leaf *E* than ambient CO_2_ equivalent plants (**Fig. 2a**). This resulted in highly significant interactions for leaf *E* dependent on CO_2_ and VPD treatment (see T2, T3 and T4 fertiliser groupings: Two-way ANOVA, *p* < 0.001; **Fig. 2a**). Previously, and in data presented here, we have shown that higher N fertiliser application enhances leaf *E* and *g_sw_* on both the abaxial and adaxial leaf surfaces simultaneously^25^ (Extended Data Fig. 1, **Fig. 2**). This response did not occur during high-VPD heatwaves for either CO_2_ treatment, with stomatal opening and *E* responses often insensitive to increasing N-application, particularly for wheat growing under ambient CO_2_ concentration (**Fig. 2b-g**, Extended Data Fig. 2a-b). Increased leaf *E* during heatwaves was particularly evident on the abaxial surface, where *E* contribution rose from a low of 8% during non-heatwave conditions, to 43-47% during heatwaves (**Fig. 2g**, Extended Data Fig. 2c). Whilst not indicative of overall water fluxes, abaxial *g_sw_* responses often followed similar trends to *E* (Extended Data Fig. 2d-h).

**Fig. 2:**
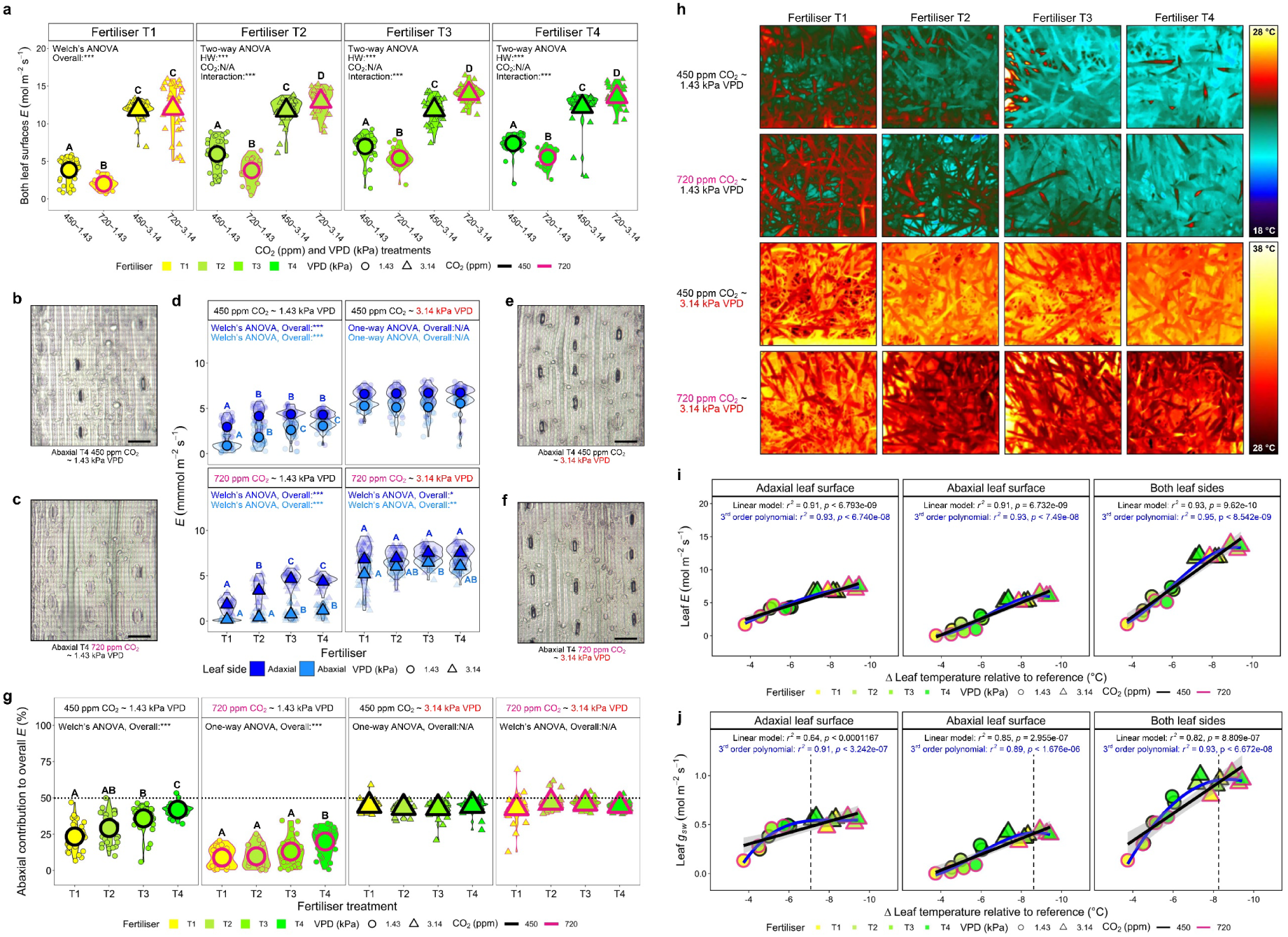
Wheat leaf and canopy responses to elevated CO_2_ and high-VPD heatwave treatments. (**a**) Porometer measured total leaf transpiration (*E*) across different CO_2_, vapour pressure deficit (VPD) and N-fertiliser treatments, measuring both leaf surfaces 5 days into heatwaves (or 5 days after flag leaves emerged during non-heatwaves). (**b-f**) Adaxial and abaxial leaf *E* of 4 CO2/VPD treatments with representative abaxial leaf surface images. (**g**) Abaxial leaf percentage contribution to total *E* under different CO_2_ and VPD. (**h**) Thermal images of wheat growing under different CO_2_, VPD and N-fertiliser treatments. Regressions of average leaf (**i**) *E* or (**j**) *g_sw_* with average Δ leaf temperature, considering adaxial, abaxial or both leaf surfaces together. For a, d, g, i-j, *n* = 32. Large symbols equal means. For One-way and Two-way ANOVAs, Tukey post-hoc tests were performed. For Welch’s ANOVAs, Games-Howell post-hoc tests were undertaken. Different letters within graphs indicate significant differences of *p ≤* 0.05. Asterisks equal, * = *p* < 0.05, ** = *p <* 0.01 and *** = *p* < 0.001.

Thermal imaging of wheat showed that high CO_2_ plants exposed to a high-VPD heatwave were typically cooler than ambient CO_2_ equivalent plants across T2, T3 and T4 treatments; validating porometry *E* measurements (**Fig. 2a-h**, Extended Data Figs. 2-3). Differences in Δ leaf temperatures measured under non-heatwave conditions also supported leaf-level *E* measurements, with highly N-fertilised plants having cooler leaves (**Fig. 2h-i**). N-fertiliser driven increases in evaporative cooling did not occur for ambient CO_2_ heatwave plants, but during high CO_2_ heatwave treatment, there was some evidence of enhanced cooling (**Fig. 2h-i**, Extended Data Fig. 3a-b). When relationships between adaxial and/or abaxial leaf *E* and Δ leaf temperature were assessed across treatments, a strong positive correlation was detected for individual leaf sides when regressed against Δ leaf temperature (both *r* ^2^ *=* 0.91, *p <* 0.001), and an even stronger relationship was detectable when both leaf side *E* values were combined (*r* ^2^ *=* 0.93, *p <* 0.001; **Fig. 2i**). Corresponding *g_sw_* measurements revealed that maximum values of adaxial leaf *g_sw_* occurred when Δ leaf temperature reached approximately −7 °C, whereas for the abaxial surface, maximum *g_sw_* was closer to −9 °C (**Fig. 2j**). These responses highlight the additive effect of abaxial *g_sw_* on plant cooling at higher VPD. Assessment of water application over the first 5 days of the heatwave revealed that more water was required by high CO_2_ heatwave plants than ambient CO_2_ equivalents, in-line with our increased *E* measurements (Two-way ANOVA, Tukey HSD, *p* < 0.001; Extended Data Fig. 3c).

### Water flux is prioritised at higher VPD

To assess the photosynthetic potential of wheat grown under current and future CO_2_/VPD scenarios, we collected leaf chlorophyll samples and measured saturating light gaseous exchanges (**Fig. 3**, Extended Data Figs. 4-5). High CO_2_, non-heatwave conditions resulted in the lowest chlorophyll content, whereas high-VPD heatwave treatment reversed this response, with high CO_2_ plants producing equal or more chlorophyll than ambient CO_2_ heatwave plants (**Fig. 3a**). Hyperspectral data analysis further confirmed the observed phenotypes, with the MERIS terrestrial chlorophyll index (MTCI) vegetation index displaying strong correlations with chlorophyll content across different CO_2_, VPD and N-fertiliser treatment combinations (Extended Data Fig. 4b).

**Fig. 3.**
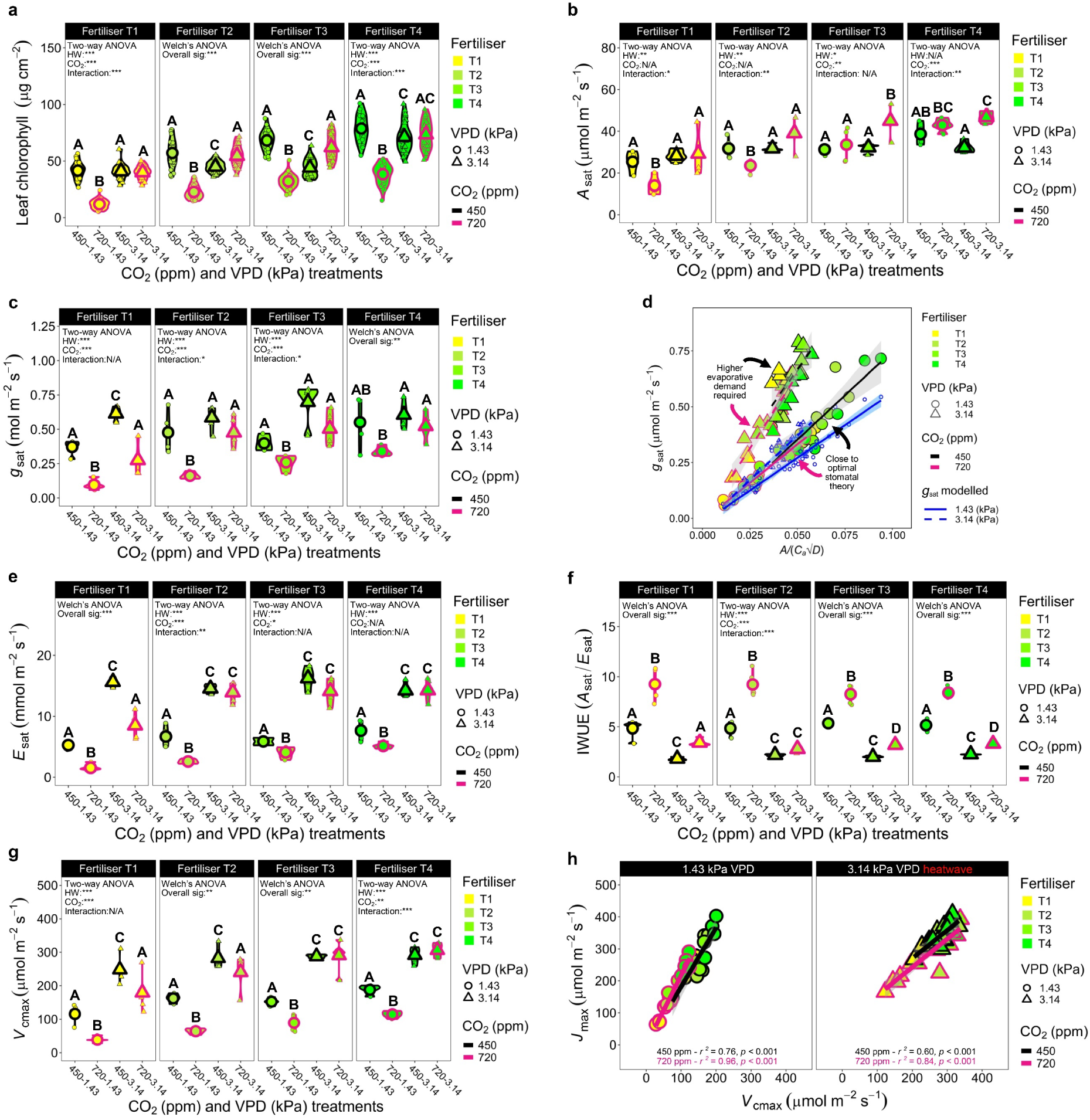
Current and future high-VPD heatwave impacts on wheat chlorophyll production and saturating light gas exchange. (**a**) Leaf chlorophyll content of plants growing under different CO_2_ concentration, vapour pressure deficit (VPD) and N-fertiliser treatments. (**b**) Photosynthesis (*A*_sat_) and (**c**) stomatal conductance (*g*_sat_), utilising infra-red gas analyser conditions that reflected CO_2_ and VPD growth conditions. (**d**) Optimal stomatal theory modelling of dependent variable *g*_sat_ with covariate *A*/(*C_a_*√*D*). Solid blue line and circle symbols represent modelled optimal *g*_sat_ at 1.43 kPa VPD (non-heatwave) and dashed blue lines and triangle symbols represents optimal *g*_sat_ at 3.14 kPa (heatwave). Black (450 ppm) and pink (720 ppm) solid and dotted lines and symbols represent equivalent regression analysis and actual values. (**e**) Saturating light transpiration (*E*) and (**f**) instantaneous water-use efficiency (IWUE). (**g**) Maximum rate of rubisco carboxylation (*V*_cmax_) and (**h**) regression analysis of *V*_cmax_ and maximum rate of electron transport (*J*_max_). For a: *n* = 32, for c-h *n* = 4-5. Except in **d**, larger symbols equal sample means. For Two-way ANOVAs, Tukey post-hoc tests were performed. For Welch’s ANOVAs, Games-Howell tests were undertaken. Different letters indicate significant differences of *p ≤* 0.05. Asterisks equal, * = *p* < 0.05, ** = *p <* 0.01 and *** = *p* < 0.001.

Higher CO_2_ increased *A*_sat_, but only for high N-fertilised plants and a consistent reduction in *g*_sat_ was also observed, but only during non-heatwave conditions (**Fig. 3b-c**). During high-VPD heatwaves, only T1 high CO_2_ plants showed significantly reduced *g*_sat_ comparatively to ambient CO_2_ equivalent plants (Two-way ANOVA, Tukey HSD, *p* < 0.001), and there were no differences between T2-T4 N-fertiliser plants. This led to significant interactions arising between CO_2_ and heatwave treatments for 3 N-fertiliser treatments (**Fig. 3c**). To further understand these responses, we modelled gaseous exchanges using optimisation theory^34,35^, which posits that stomata operate to maximise *A* and minimise transpirational water loss (**Fig. 3d**). Measured *g*_sat_ values were considerably higher than predicted by the model during high-VPD heatwaves, with the slopes of measured ambient and high CO_2_ plants not dissimilar (Welch’s t-test, test statistic = 0.35, *p* = 0.73). This reveals that stomatal behaviour changed markedly to preferentially favour water release over photosynthetic gain during heatwaves, irrespective of CO_2_ concentration. These altered gas exchange strategies were linked with significantly higher leaf VPD values that also corresponded with higher overall *C*_i_:*C*_a_ ratios (Extended Data Fig. 4e-f).

Further assessment of gas exchange water fluxes revealed non-heatwave, high CO_2_ plants had the lowest saturating light transpiration (*E*_sat_), whereas heatwave imposition routinely led to much higher *E*_sat_, with no significant differences detectable between CO_2_ treatments, except for T1 plants (**Fig. 3e**, Extended Data Fig. 4g). High CO_2_ wheat routinely had higher IWUE (*A*/*E*) and iWUE (*A*/*g_sw_*) within each VPD treatment, (**Fig. 3f**, Extended Data Fig. 4h), although because *E*/*g_sw_* values were much higher during high-VPD heatwaves, the IWUE and iWUE of high CO_2_ plants were markedly lower than during non-heatwaves. To assess whether CO_2_ and/or heatwaves impacted photosynthetic biochemistry, we investigated the maximum rate of rubisco carboxylation (*V*_cmax_) and maximum rate of photosynthetic electron transport (*J*_max_) (**Fig. 3g**, Extended Data Fig. 5a). Heatwaves boosted *V*_cmax_ (especially for highly fertilised wheat), whereas *J*_max_ values often remained more similar. This led to obvious slope differences between ambient VPD and heatwave scenarios when *J*_max_ and *V*_cmax_ were regressed (**Fig. 3h**). *A*/*C*_i_ curve analysis revealed high CO_2_ plants were mostly RuBP limited under growth CO_2_ concentration, whereas ambient CO_2_ plants were typically rubisco limited. Supply function comparisons between ambient and high CO_2_ treatments revealed that stomatal limitations were the same between highly fertilised plants experiencing high-VPD heatwave conditions, but this was not the case under lower N-fertiliser treatments and/or ambient VPD (Extended Data Fig. 5b-i).

### High VPD inhibits stomatal dynamism

Enhancing stomatal responsiveness during changing environmental conditions is a key focus for boosting crop iWUE and *A*^36^. Light-shift experiments (saturating light steady-state: 5 mins → dark: 1hr → saturating light: 1hr) were undertaken on high N-fertilised, T4 wheat plants under conditions mimicking the growth environments of the 4 different CO_2_/VPD scenarios (**Fig. 1 and 4**). For non-heatwave treatments, high CO_2_ plants initially had similar *A* to ambient CO_2_ plants, whereas *g_sw_* was significantly lower (Two-way ANOVA, *p* < 0.01; **Fig. 4a**, Extended Data Fig. 6a-b). For high-VPD heatwaves, ambient CO_2_ plants started with significantly lower *A* than high CO_2_ equivalent plants (Two-way ANOVA, *p* < 0.001), whereas *g_sw_* values was not significantly different (**Fig. 4b**, Extended Data Fig. 6a-b). Upon dark application during non-heatwaves, ambient CO_2_ plants took much longer for stomata to close (∼60 mins relative to ∼20 minutes for high CO_2_), but overall, the initial rate of stomatal closure during the first 20 minutes of dark was not significantly different (**Fig. 4a and c**). Heatwaves considerably hampered stomatal closing for both CO_2_ treatments, with *g_sw_* values only reducing by approx. 37-38%, and no rate change differences were detectable between ambient and high CO_2_ plants (**Fig. 4b-d**).

**Fig. 4:**
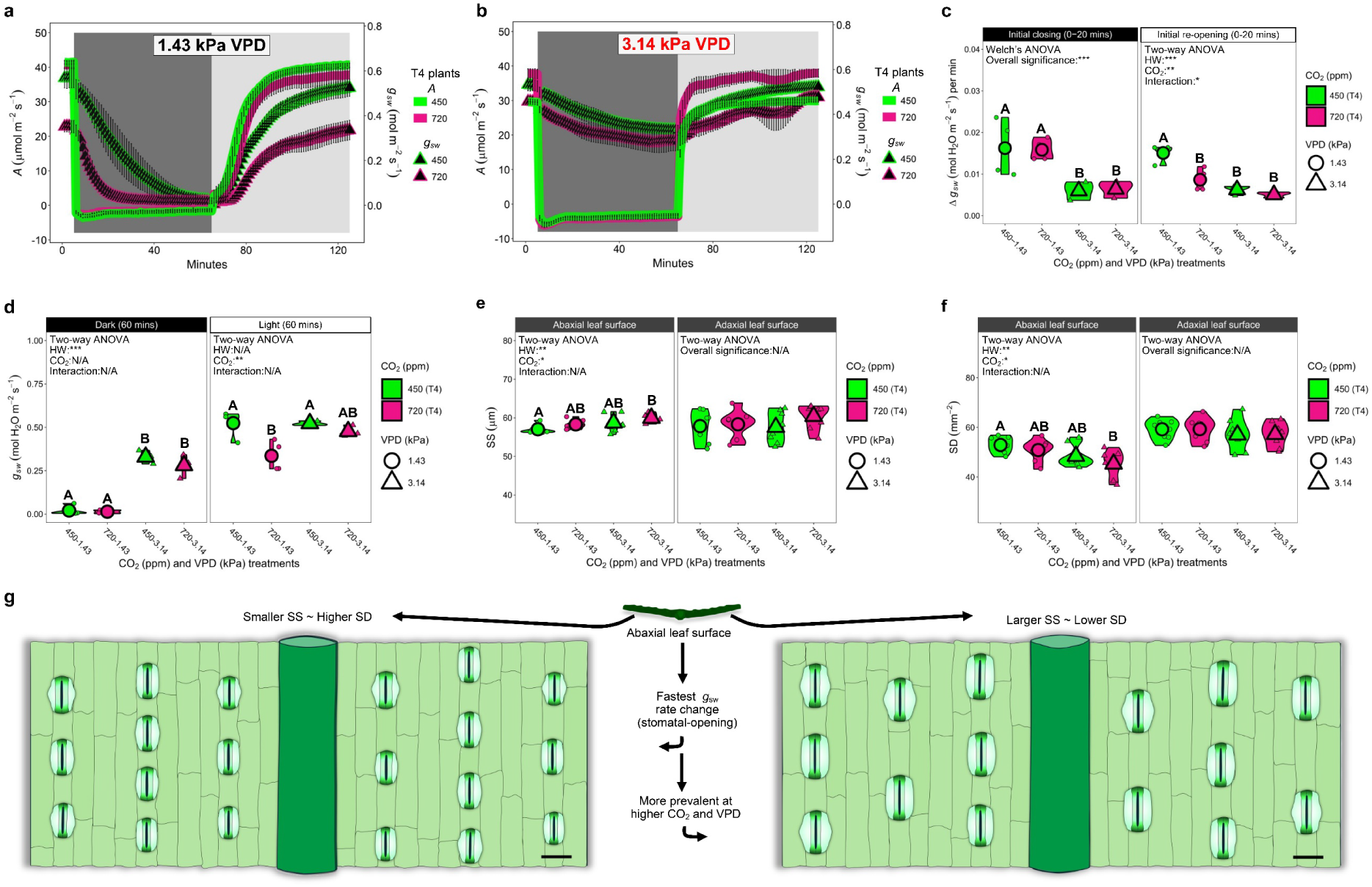
Wheat stomatal dynamics during different CO_2_ and VPD growth conditions. (**a**-**b**) Light-saturated light-shift experiments employing growth chamber CO_2_ concentrations and VPD settings. Gaseous exchanges were recorded every minute for 125 minutes, with 5 minutes saturating light then 1 hr of dark measurements, and finally a further 1 hr of saturating light measurements. Results plotted separately for (**a**) non-heatwave (VPD: 1.43 kPa) and (**b**) heatwave (VPD: 3.14 kPa) treated plants. (**c**) Initial *g_sw_* rate changes during closure period (20 minutes) and re-opening period (20 minutes) for a-b. (**d**) Measurements of *g_sw_* at the end of the dark period (65 minutes) and the end of light re-application (125 minutes). Abaxial and adaxial (**e**) stomatal size (SS) and (**f**) stomatal density (SD) measurements. (**g**) SS and SD alterations in response to different CO_2_∼VPD conditions. For a-d: *n* = 3-4, for e-f *n* = 8. Large symbols equal sample means. See graphs for statistical tests. For Two-way ANOVAs, Tukey post-hoc tests were performed. For Welch’s ANOVAs, Games-Howell tests were undertaken. Different letters indicate significant differences of *p ≤* 0.05. Asterisks equal, * = *p* < 0.05, ** = *p <* 0.01 and *** = *p* < 0.001.

Upon saturating light re-application, ambient CO_2_, non-heatwave plants had a faster *g_sw_* rate change than high CO_2_ equivalent plants (**Fig. 4a, c**). Heatwave treatments led to the slowest *g_sw_* rate changes, with no difference between ambient and high CO_2_ plants (**Fig. 4b-c**). After 60 minutes of saturating light, ambient CO_2_ plants again had significantly higher *g_sw_* than high CO_2_ plants (during non-heatwaves), but this was not the case for heatwave-treated plants, where *g_sw_* values between CO_2_ treatments were similar (**Fig. 4d**). To further evaluate stomatal responsiveness to higher VPD, we also assessed *E* and *g_sw_* values from *A*/*C*_i_ curves of T4 plants and found that heatwave treatment led to CO_2_ blindness at higher CO_2_ reference concentrations irrespective of CO_2_ growth concentration, with no sign of stomatal closure (Extended Data Fig. 7).

Stomatal size (SS) and stomatal density (SD) assessment on both leaf surfaces revealed the abaxial leaf surface to be more responsive to different growth conditions, whereas the adaxial surface did not show such plasticity (**Fig. 4e-g**). Overall, both CO2 and VPD affected abaxial SS, with higher CO2 and higher VPD increasing SS (Two-way ANOVA, CO_2_ = *p* < 0.05, HW = *p* < 0.01; **Fig. 4e**). Increased VPD also significantly reduced abaxial SD (Two-way ANOVA, *p* < 0.01), and the combination of both high CO2 and VPD led to significant reductions in SD comparatively to ambient CO_2_ and VPD plants (Two-way ANOVA, Tukey HSD, *p* < 0.01; **Fig. 4f**).

### Heatwaves diminish CO_2_ drought responses

Removing irrigation revealed marked differences in drought stress resilience, dependent on CO_2_ and VPD treatment (**Figs. 5 and 6**). Non-heatwave, high CO_2_ plants often performed best under drought conditions, with 2 out of 4 N-fertiliser treatments (T2 and T4) maintaining significantly higher *E* than all other growth scenario plants at day 6 (Generalised linear model (GLM), *p* ≤ 0.01; **Fig. 5a-g**). MTCI assessment, which serves as a proxy for leaf chlorophyll content (Extended Data Figs. 1 and 4), also confirmed non-heatwave, high CO_2_ plants as best performing, with green leaves still often visible at day 6 (GLM, T2:T4: *p* ≤ 0.01; **Fig. 5a-d, h-j**). For ambient and high CO_2_ high-VPD heatwave plants, *E* and MTCI values did not show such clear differences at day 6. By drought day 12, water was mainly absent from the upper portions of T4 plant canopies due to desiccation, except for non-heatwave, high CO_2_ plants, which still displayed green ears. This led to significantly warmer ear temperatures comparatively to the dried-out white ears from the other 3 VPD∼CO_2_ treatments, which reflected a greater proportion of incoming radiation and were thus cooler (Two-way ANOVA, Tukey HSD, *p* ≤ 0.01, **Fig. 5a-d, k-l**).

**Fig. 5:**
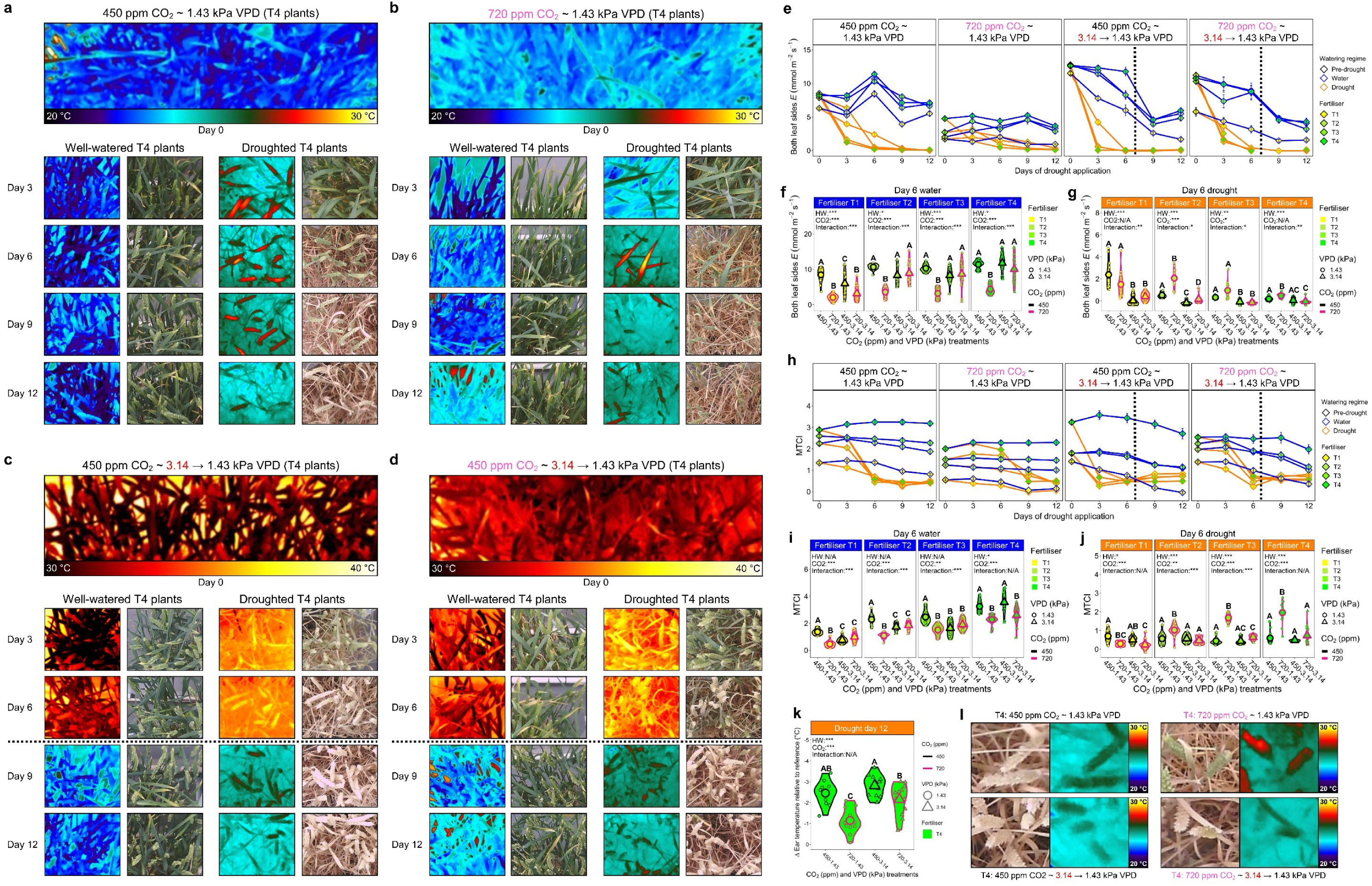
Wheat drought resilience under current and future high-VPD heatwave conditions. (**a**-**d**) Thermal and digital images of T4 wheat plants during (**a**) 450 ppm CO_2_, non-heatwave (VPD: 1.43 kPa), (**b**) 720 ppm CO_2_, non-heatwave, (**c**) 450 ppm CO_2_, heatwave (VPD 3.14 kPa) and (**d**) 720 ppm CO_2_ heatwave conditions. At day 7, VPD was reduced from 3.14 kPa → 1.43 kPa for heatwave plants (see dotted lines). (**e**) Combined leaf surface *E* during drought. (**f**-**g**) Day 6 quantification of *E* values for (**f**) continually watered plants and (**g**) droughted plants. (**h**) MERIS terrestrial chlorophyll index (MTCI) responses during drought. (**i**-**j**) Day 6 MTCI of (**i**) continually watered plants and (**j**) droughted plants. (**k**) Droughted wheat ear temperatures at day 12 of drought. (**l**) Thermal and digital images of wheat ears. For e-j: *n* = 32, for k: *n* = 8. Large symbols equal sample means. Generalised linear models with post-hoc assessment via estimated marginal means were undertaken for significance. Different letters indicate significant differences of *p ≤* 0.05. Asterisks equal, * = *p* < 0.05, ** = *p <* 0.01 and *** = *p* < 0.001.

**Fig. 6:**
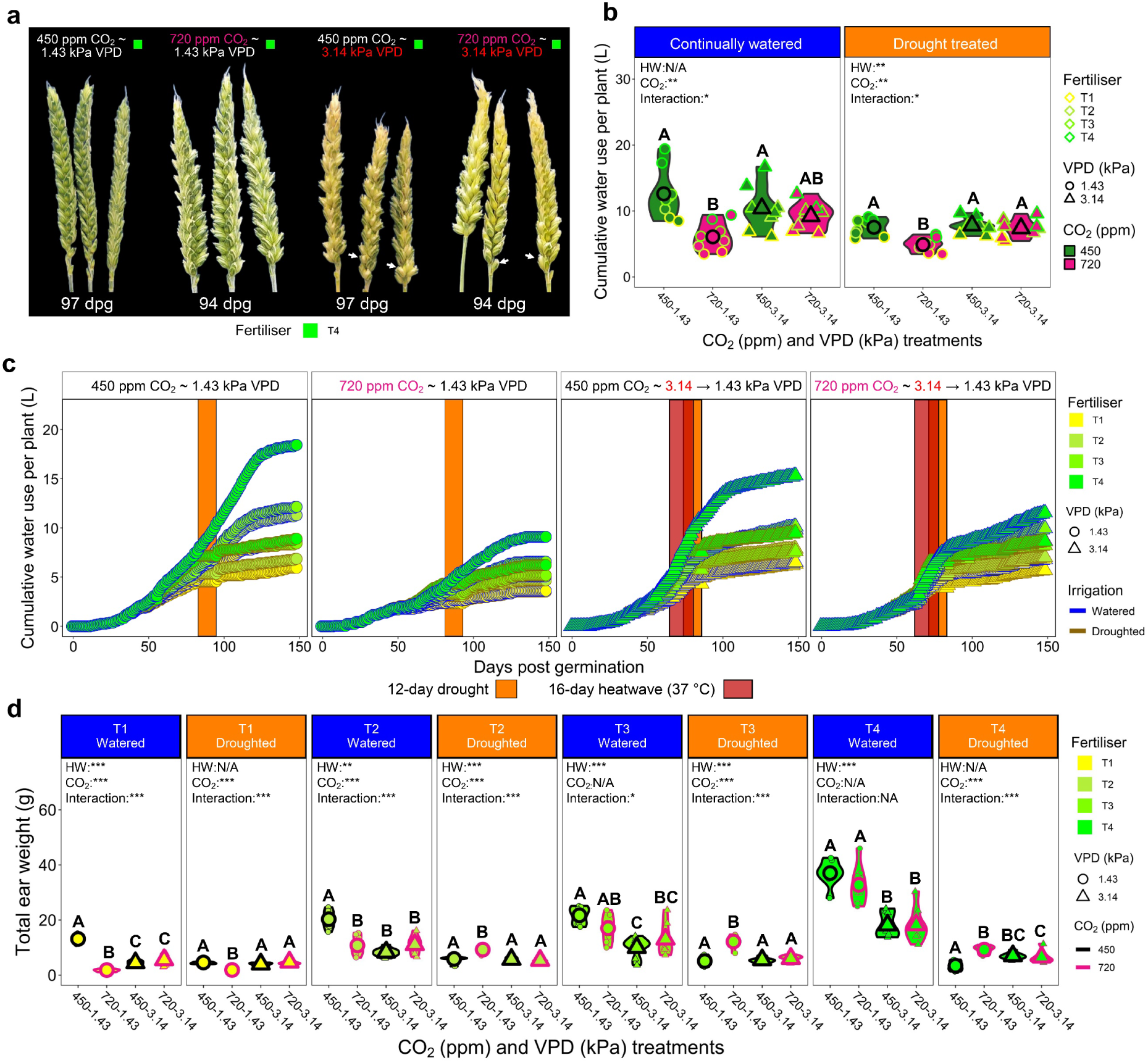
The impacts of high CO_2_, heatwaves and N-fertiliser application on wheat seasonal water usage and productivity. (**a**) RBG images of wheat ears post-heatwave and drought treatment. * Note example bulbous grains (white arrows) developed on high-VPD heatwave ears, but not on non-heatwave ears. (**b**) Quantified total water usage and (**c**) water usage throughout 148-day growing season for 4 different VPD∼CO_2_ treatment combinations. (**d**) Total ear weight of well-watered and droughted plants. For b and d, *n* = 8, for c, *n* = 2. Larger symbols equal sample means. Statistical testing undertaken using generalised linear models with post-hoc assessment conducted by computing estimated marginal means. Different letters within graphs indicate significant differences of *p ≤* 0.05. Asterisks equal, * = *p* < 0.05, ** = *p <* 0.01 and *** = *p* < 0.001.

Heatwaves increased the speed of wheat development, which led to faster declines in MTCI values (**Fig. 5h**). This quicker maturation was also seen during ear and grain development, with ears that developed under heatwaves quicker to develop bulbous grains (**Fig. 6a**). Assessment of total water application over the 148-day duration of experiments, confirmed non-heatwave, high CO_2_ plants used approx. 52% less water per plant (6.09 L) than ambient CO_2_ equivalent plants (12.6 L) when continually watered (GLM, *p* < 0.001; **Fig. 6b-c**). There were no significant differences in water application between CO_2_ treatments during well-watered or droughted high-VPD heatwave conditions, but continually watered T4 ambient CO_2_ wheat did exhibit a trend towards receiving more water than high CO_2_ equivalent heatwave plants (Two-way ANOVA, Tukey HSD, *p* = 0.19; **Fig. 6b-c**).

N-Fertiliser was a crucial determinant of total above ground biomass and ear weight (a proxy for yield, **Fig. 6c**, Extended Data Fig. 8). Non-heatwave, high CO_2_ plants almost always produced the highest total ear weight during drought (T2-T4 N-fertiliser treatments), but this was not the case for total above ground biomass accumulation (**Fig. 6c**, Extended Data Fig. 8). High-VPD heatwaves impacted ear weight and total biomass most in highly fertilised plants, and this typically occurred irrespective of CO_2_ growth conditions. Assessment of water-use efficiency (WUE; total ear weight/ total water applied) over the whole lifecycle revealed highly fertilised T4 wheat grown under non-heatwave, high CO_2_ conditions had the highest WUE, requiring 286 ml for every gram of ear, comparatively to 501 ml for ambient CO_2_ plants (GLM, *p* < 0.01; Extended Data Fig. 8). Whilst high-VPD heatwaves reduced overall WUE, we did detect higher WUE in T4, high CO_2_ plants (676 ml) than ambient CO_2_ equivalents (866 ml) (GLM; 0.05; Extended Data Fig. 8).

## Discussion

A robust understanding of how cultivated crops respond to environmental extremes will be crucial for developing future climate-resilience. Wall et al.^37^, previously highlighted the dominant effect of the wheat adaxial leaf surface on gaseous exchanges, showing adaxial leaf *A* and *g_sw_* contributions to be approximately twice that of abaxial leaf surfaces. Here, by assaying both wheat leaf surfaces within actively growing canopies, we highlight how abaxial and adaxial leaf surface contributions to *g_sw_* and *E* depend on multiple factors including: the concentration of atmospheric CO_2_, N-fertiliser quantity and the prevailing VPD growth conditions (**Fig. 2**). Higher CO_2_ concentrations typically restrict *E*/*g_sw_* under ambient VPD, whereas higher N-Fertiliser has the opposite response. During high-VPD heatwaves, both CO_2_ and N-Fertiliser stomatal responses become attenuated, with higher evaporative water release from abaxial leaf surfaces increasingly important (**Figs 2-4**). Overall, VPD-driven stomatal blindness to rising CO_2_ coupled with much higher photosynthetic capacity, diminished the positive drought responses often associated with higher CO_2_ concentration. This led to droughted wheat ear weights of high CO_2_ and ambient CO_2_ heatwave plants not being significantly different (**Figs. 5-6**).

There is limited research covering the combined effects of CO_2_, VPD, temperature and N deposition on crop gaseous exchanges and productivity, and so climate models are currently limited with regard parametrisations that intertwine these 4 key growth factors. High CO_2_ or high VPD often reduce *g_sw_*^38–40^, whereas the interactive effects of both factors has been shown to reduce such decreases^41,42^ and recent global meta-analysis suggests that the combined effects of high N-deposition and rising temperature oppose *g_sw_* reductions^43^. Here, we show during heatwave treatments (with moderately high VPD at high temperature), that stomatal light and CO_2_ responses of *g_sw_* are diminished (or absent) in high N-fertilised plants, and this occurs irrespective of CO_2_ growth concentration. Such responses prompt a move away from optimal stomatal behaviour^34^ to favour increased water release, and for high CO_2_ plants, this may well have been linked with greater nutrient requirements to support higher rates of *A*, with chlorophyll production clearly upregulated in response to high VPD and high CO_2_ **(Fig. 3b**). Despite lower measured *A_sat_* values, ambient CO_2_ heatwave plants maintained similar *g_sat_* and *E* to high CO_2_ equivalent plants, which implies decoupling was occurring as higher water fluxes were still essential. Given the substantial reductions in ear weight of all heatwave plants, we suggest that this maintained high water release may well have been an attempt to mitigate against the impacts of temperature-inducted infertility during flowering, which is known to reduce grain yields by approx. 3-5% for every 1 degree above optimum temperatures^44^.

VPD-driven differences in stomatal responsiveness occurred alongside differences in stomatal SS and SD, with heatwave imposition and higher CO_2_ both contributing to larger SS and smaller SD, but only on abaxial leaf surfaces (**Fig. 4**). SS∼SD responses are also often detected on fossilised leaves of plants that grew under high CO_2_ climates (larger SS, lower SD)^45^, which leads to lower calculated maximum stomatal conductance (*g*_smax_). Natural rice varieties with larger SS and smaller SD, have also been shown to display lower *g*_smax_, with corresponding plants having higher *E* (without reducing *g_sw_*) when exposed to rising VPD (experiments at ambient CO_2_ concentration)^8^. Smaller stomata are suggested to enable quicker responsiveness to environmental stimuli due to greater surface area to volume ratios^46–48^ and coupled with higher SD, this has been suggested to boost carbon uptake via higher *g*_smax_ when atmospheric CO_2_ concentrations were low^45^. However, if the need to release water at moderately higher VPD overrides requirements to close stomata in presence of low light (or higher CO_2_), then having overly responsive stomata might hinder plant performance. If this is the case, then optimising stomatal *g_sw_*, via smaller SS and larger SD (for enhanced iWUE), might not be tenable for future climate extremes.

Taken together our results show that wheat stomata are insensitive to CO_2_-induced stomatal closure during high-VPD heatwaves. This leads to high CO_2_ grown plants using considerably more water than ambient CO_2_ equivalent plants during the initial stages of heatwaves, with increased evaporative demand potentially linked to higher photosynthetic capacity. Stomatal adjustments in response to irradiance changes (and rising CO_2_) were also significantly impaired, which together suggests that future wheat could be considerably less water-use efficient than previously envisioned. Research is now urgently required to understand how high VPD heatwaves and rising CO_2_ impact stomatal-regulated water losses in other plant species, so that global modelling efforts and crop resilience can be optimised for future climates.

## Methods

### Plant materials and growth conditions

Experiments were conducted on the elite British wheat (*Triticum aestivum L*) variety Mulika (Blackman Agriculture Ltd, Cambridge, UK). Individual plants were grown in 0.8 l pots (IPP, Bytom, Poland) with 5:1 Levington M3: Perlite compost mix. Chamber CO_2_ concentrations were set to 450 ppm or 720 ppm CO_2_ for the entirety of experiments with plants gradually acclimatised to peak growth conditions over the first 4 weeks of seed germination. For the first 2 weeks, chambers were set to 15 °C day: 10 °C night, with a 12h: 12h, light: dark cycle with photosynthetically active radiation (PAR) of 1000 μmol m^−2^s^−1^ provided at canopy level. This relatively high light treatment increased temperatures around the canopy by +4 °C, so growth temperatures around plants were approx. 19 °C during the day. Over the next two weeks, chamber set temperatures were gradually increased to 23 °C during the day and 15 °C during the night, with daylength gradually increased from 12 to 14 hours. Because of the high light intensity, the temperature at canopy level was approx. 27 °C by week 4. Relative humidity was set continually at 60% throughout experiments except during heatwaves where it was 50%. For the 16 days during heatwave treatments, chamber growth temperatures were increased by 10 °C day and night (33 °C day: 25 °C night), which meant around plant canopies it was approximately 37 °C during the day. Post-heatwaves, cabinet growth temperatures were returned to previous levels.

A total of 128 plants were grown for each of the 4 growth scenarios, with deionised water (dH_2_O) applied to pot bases in trays when required. From day 32 of each growth scenario, plants were divided into 4 fertiliser groupings (32 plants for each fertiliser treatment: T1-T4) and if applicable, were fed with high nitrogen fertiliser on a weekly basis for 12 weeks (Chempak® High Nitrogen Feed - Formula 2, Thompson and Morgan, Southampton, UK). T1 plants were no fertiliser controls, T2 plants received 2.125g, T3 plants received 4.25g and T4 plants received 8.5g. Soluble fertiliser was dissolved in 2 litres of water and divided equally between 4 trays that held the 32 plants of each fertiliser grouping. Droughted plants that missed fertiliser treatment received additional applications at weekly intervals after the 12 initial applications, so that all fertiliser treatment plants received a total of 12 applications.

### Plant gas exchange measurements

Leaf stomatal conductance to water vapour (*g_sw_*) and transpiration (*E*) were collected using a LI-600 porometer (LI-COR, Lincoln, OR, USA) set to a flow rate of 150 μmol s^−1^. Saturating light steady-state gas exchange was collected using a LI-COR LI-6800 Portable Photosynthesis Systems with attached Multiphase Flash Fluorometer (6800-01A). The leaf chamber conditions were set according to conditions at the canopy, with the exception of light intensity which was set to a saturating light value of 1800 μmol m^−2^ s^−1^ PAR. The flow rate was 400 μmol s^−1^, with CO_2_ concentration controlled by CO_2_ reference and temperature by Tair. For each plant, 10 readings were taken under steady-state conditions over a 5-minute period and averaged (*n* = 4-5 plants). For saturating light stomatal response curves, 5-minutes of steady state measurements were first collected, before irradiance was removed for 1 hour and then reapplied for an hour. Like steady-state measurements, conditions apart from light matched the surrounding canopy.

To produce *A*/*C*_i_ curves to model photosynthetic traits, the same leaf chamber conditions were used as above for each growth scenario, except T_leaf_ was set to either 27°C or 37°C. The [CO_2_]_ref_ sequence applied for 450 ppm plants was: 450, 325, 200, 150, 100, 75, 50, 25, 450, 450, 450, 450, 585, 720, 860, 1000, 1250, 1500, 1800. For plants grown at 720 ppm the order of [CO_2_]_ref_ was 720, 585, 450, 325, 200, 150, 100, 75, 50, 25, 720, 720, 720, 720, 860, 1000, 1250, 1500, 1800 ppm. A match was conducted prior to each measurement with 3-5 minutes permitted between each [CO_2_]_ref_ treatment for stabilisation. Steady-state, *A*/*C*_i_ and light response curves were collected between days 4-9 of heatwave treatments. The maximum carboxylation rate of rubisco (*V_cmax_*) and the maximum rate of photosynthetic electron transport (*J_max_*) were modelled using the Plantecophys package on R software^49^. For leaf stomatal analysis methodology see Caine et al.,^8^.

### Thermal imaging

Thermal images of plants were captured using a FLIR T540 thermal imaging camera (Wilsonville, OR, USA). One leaf from each plant was measured and subtracted from the temperature of the green plastic hemispherical dry reference surface to calculate relative leaf temperature differences across the different growth scenarios. Images analysis was conducted in FLIR Researcher IR MAX (www.flir.co.uk).

### Hyperspectral imaging and data analysis

A PSR+ spectroradiometer (Spectral Evolution, Haverhill, MA, USA) was used to collect leaf-level hyperspectral reflectance data (350-2500 nm), utilising an attached Spectral Evolution leaf clip assembly. Target and reference panel measurements from a radiometrically calibrated 99% Spectralon panel were sampled sequentially. The Spectrolab package was used to import hyperspectral data in R^50^.

### Leaf biochemical and structural measurements

For analysis of leaf chlorophyll content, 2x 6 mm diameter leaf discs were collect at day 8 of the heatwave and then weighed and placed into 5 ml of N,N-Dimethylformamide ≥99.8% ACS (Thermo scientific, Oxford, UK) and stored at 4 °C for 7 days to enable chlorophyll extraction. Chlorophyll absorbance was measured at following wavelengths: 663.8, 646.8 and 480 nm^51^, with a Shimadzu UV2600i spectrophotometer (Kyoto, Japan). For leaf stomatal analysis see Caine et al.^8^.

### Stomatal optimisation modelling

The optimal stomatal theory equation for computing optimal stomatal performance for different CO_2_ and VPD growth treatments is set out in equation 1:

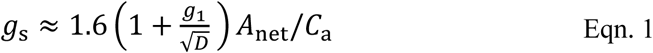

Where *g*_s_ is the stomatal conductance, *D* is the leaf–air vapour pressure difference (kPa), *A*_net_ is measured photosynthesis, and *C*_a_ is the CO_2_ concentration outside the leaf. To estimate *g*_1_, the equation used in^34^ was utilised, as set out in equation 2 where *C*_i_ refers to the CO_2_ concentration inside of the leaf:

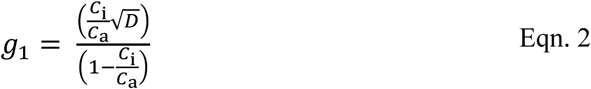

## Acknowledgements

This work was supported by the UK Research and Innovation (UKRI) Future Leaders Fellowship scheme [MR/T01993X/1] and The Institute of Sustainable Food at the University of Sheffield.

## Author contributions

RSC and HLC designed the study. RSC, MSK and YS collected the data. RSC undertook data analysis. RSC and HLC wrote the manuscript with input from MSK, YS and CPO.

## Materials & Correspondence

Please contact RSC or HLC should any data be required.

