## Supplementary data for "Future heatwave conditions inhibit CO_2_-induced stomatal closure in wheat"

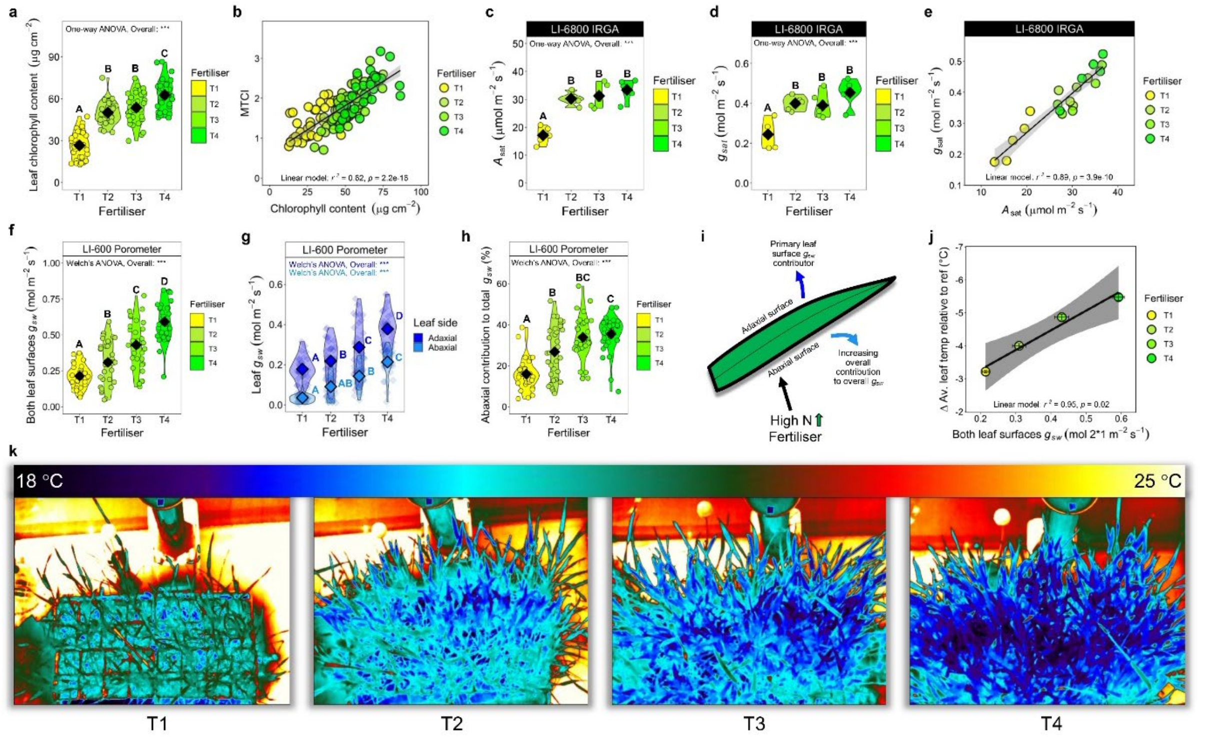


**Extended Data Fig. 1. Increased application of high-N fertiliser enhances gaseous exchanges on both leaf surfaces of cultivated wheat.** (**a**) Leaf chlorophyll content of wheat supplied with increasing concentrations of high-N fertiliser (T1 least T4 most). (**b**) MERIS terrestrial chlorophyll index (MTCI) vegetation index regressed again leaf chlorophyll content. LI-COR LI-6800 Infra-red gas analyser measurements of (**c**) saturating light photosynthesis ($A_{sat}$) and (**d**) saturating light stomatal conductance ($g_{sat}$). (**e**) Regression of $g_{sat}$ and $A_{sat}$. LICOR LI-600 porometer measurements of (**f**) total $g_{sw}$ from both leaf sides, (**g**) individual adaxial and abaxial contributions, and (**h**) abaxial % contribution to total $g_{sw}$. (**i**) Schematic representing N-fertiliser impacts on gaseous exchange. (**j**) Combined averaged canopy-level $g_{sw}$ measurements of both leaf surfaces regressed against averaged Δ leaf temperature of leaves relative to an in-chamber reference surface. (**k**) Thermal images of T1-T4 wheat canopies, with thermal scale. $N = 32$ in a, b, f, g, h and j. Large symbols equal sample means. For one-way and two-way ANOVAs, Tukey post-hoc tests were performed to determine significance. For Welch's ANOVAs, Games-Howell post-hoc tests were undertaken. Different letters within graphs indicate significant differences of $p \leq 0.05$. Asterisks equal, * = $p < 0.05$, ** = $p < 0.01$ and *** = $p < 0.001$.

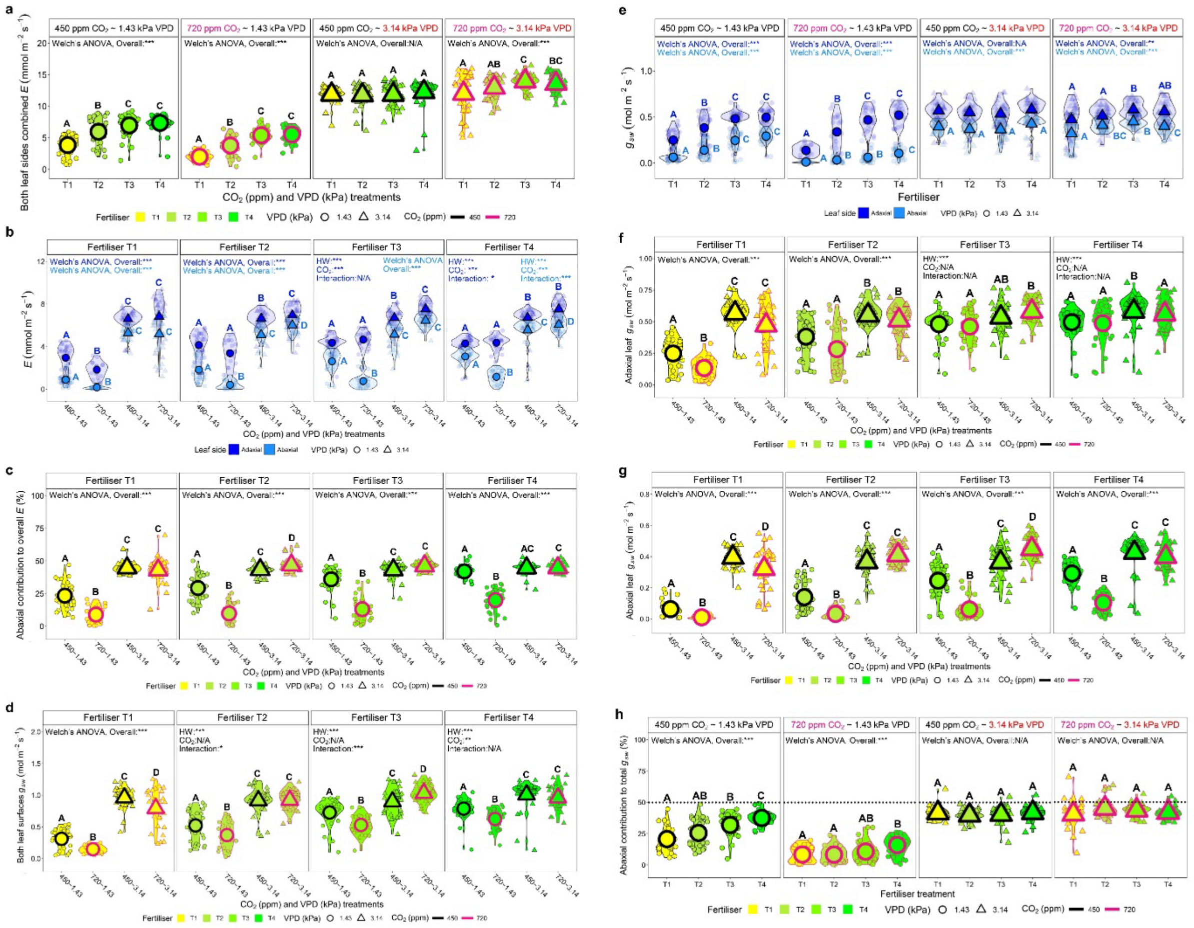


**Extended Data Fig. 2. The impacts of $CO_2$, VPD and N-fertiliser treatment on leaf gaseous exchanges.** (**a**) Porometer measurements of transpiration (*E*) of both leaf surfaces across a gradient of nitrogen fertiliser treatments, grouped via ambient (450 ppm) or high $CO_2$ (720 ppm), under non-heatwave (VPD = 1.43 kPa) or heatwave (VPD = 3.14 kPa) conditions (**b**) Abaxial and adaxial *E* leaf measurements grouped by N-fertiliser treatment across growth treatment scenarios. (**c**) Abaxial leaf surface contribution to *E* grouped by N-fertiliser treatment. (**d**) Both leaf surfaces combined stomatal conductance to water vapour ($g_{sw}$) grouped by N-fertiliser treatment. (**e**) Abaxial and adaxial $g_{sw}$ measurements across N-fertiliser treatment grouped by growth treatment scenarios. (**f**) Adaxial and (**g**) abaxial $g_{sw}$ grouped via grouped by N-fertiliser treatment. (**h**) Abaxial $g_{sw}$ contribution to total $g_{sw}$ across N-fertiliser gradient grouped by growth scenario. $n = 32$. Different letters of the same colour within graphs represent significant differences between treatments. Statistical tests equal Two-way ANOVAs unless otherwise stated within graphs, with * = $p < 0.05$, ** = $p < 0.01$ and *** = $p < 0.001$.

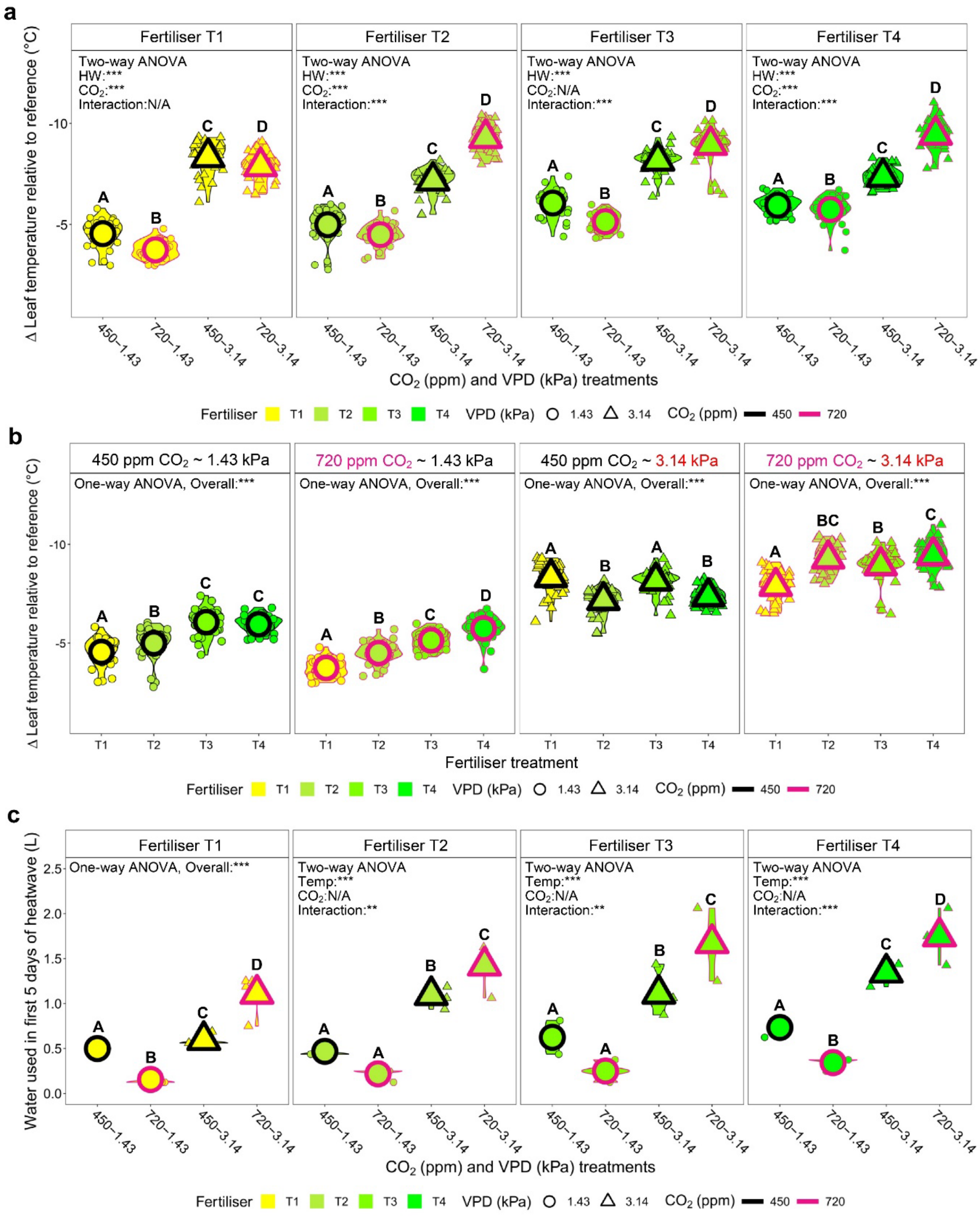


**Extended Data Fig. 3. Infrared thermography and whole-plant water application of wheat grown under different $CO_2$ and heatwave treatments.** (**a**) The Δ leaf temperature differences of different $CO_2$ and vapour pressure deficit (VPD) treated plants, grouped by N-fertiliser treatment. (**b**) The Δ leaf temperature differences of different $CO_2$ and vapour pressure deficit (VPD) treated plants, grouped by growth scenario treatment. (**c**) Water applied over the first 5 days of heatwave. For a-b: $n = 32$, for c: n = 4. Large symbols equal sample means. For One-way and Two-way ANOVAs, Tukey post-hoc tests were performed. For Welch's ANOVAs, Games-Howell post-hoc tests were undertaken. Different letters within graphs indicate significant differences of $p \leq 0.05$. Asterisks equal, * = $p < 0.05$, ** = $p < 0.01$ and *** = $p < 0.001$.

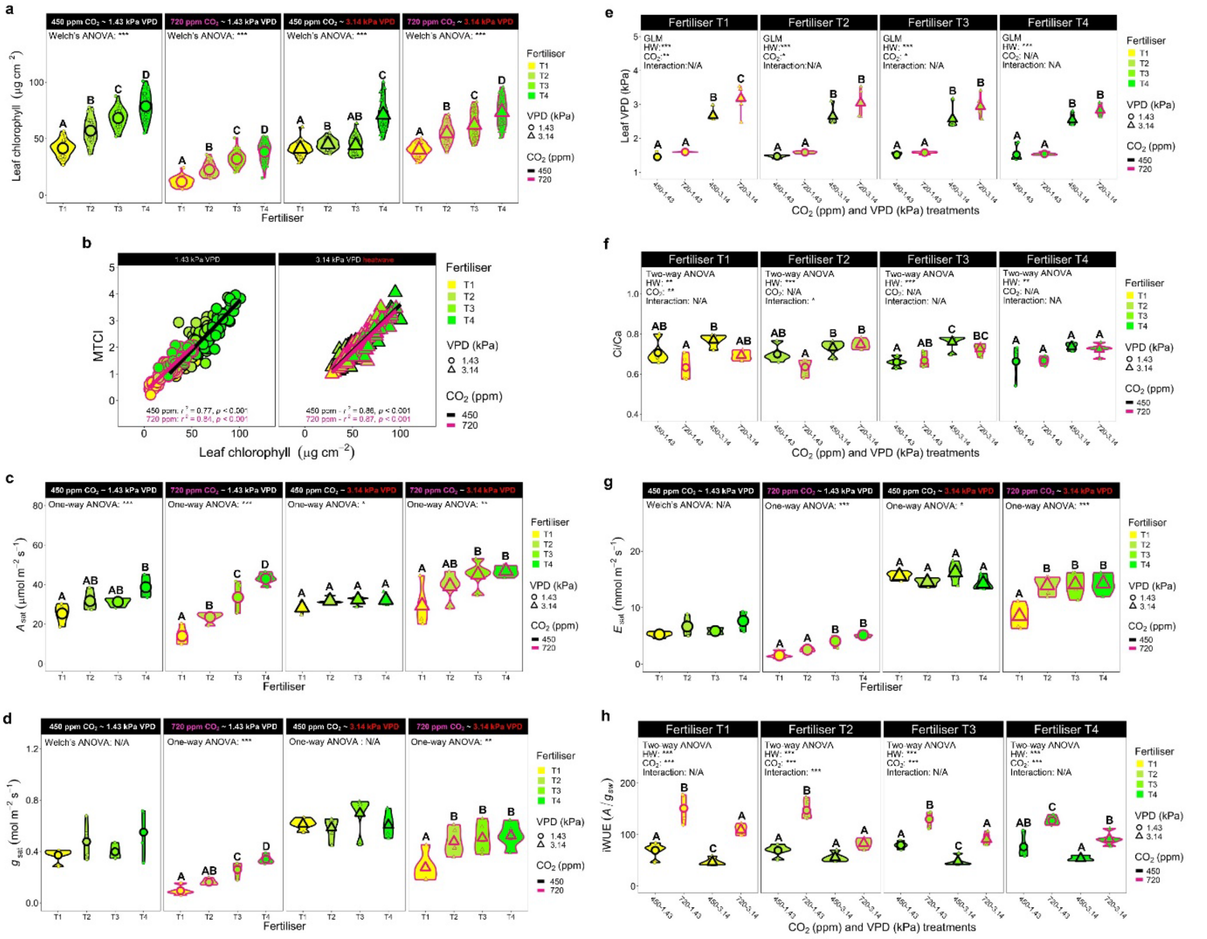


**Extended Data Fig. 4. Further chlorophyll and saturating light gas exchange analysis of plants grown under different $CO_2$ concentrations and heatwave scenarios.** (**a**) Leaf chlorophyll content of plants growing under different $CO_2$ concentration, vapour pressure deficit (VPD) and N-fertiliser treatments grouped via growth scenario. (**b**) Leaf chlorophyll content regressed against MERIS terrestrial chlorophyll index (MTCI). (**c**) Photosynthesis ($A_{sat}$) and (**d**) stomatal conductance ($g_{sat}$), utilising the same infra-red gas analyser conditions that reflected plant $CO_2$ and VPD growth treatments, grouped via growth scenario. (**e**) Leaf VPD of different growth scenario plants grouped via N-fertiliser treatment. (**f**) Equivalent Intercellular $CO_2$ concentration (Ci) divided by ambient $CO_2$ concentration, grouped via N-fertiliser treatment. (**g**) Saturating light transpiration ($E$) grouped by growth treatment scenario (**h**) Intrinsic water-use efficiency (iWUE) grouped by fertiliser group. For a and b: $n = 32$, for c-h n = 4-5. Except in **b**, large symbols equal sample means. For One and Two-way ANOVAs, Tukey post-hoc tests were performed. For Welch's ANOVAs, Games-Howell tests were undertaken. Different letters indicate significant differences of $p \leq 0.05$. Asterisks equal, $* = p < 0.05$, $** = p < 0.01$ and $*** = p < 0.001$.

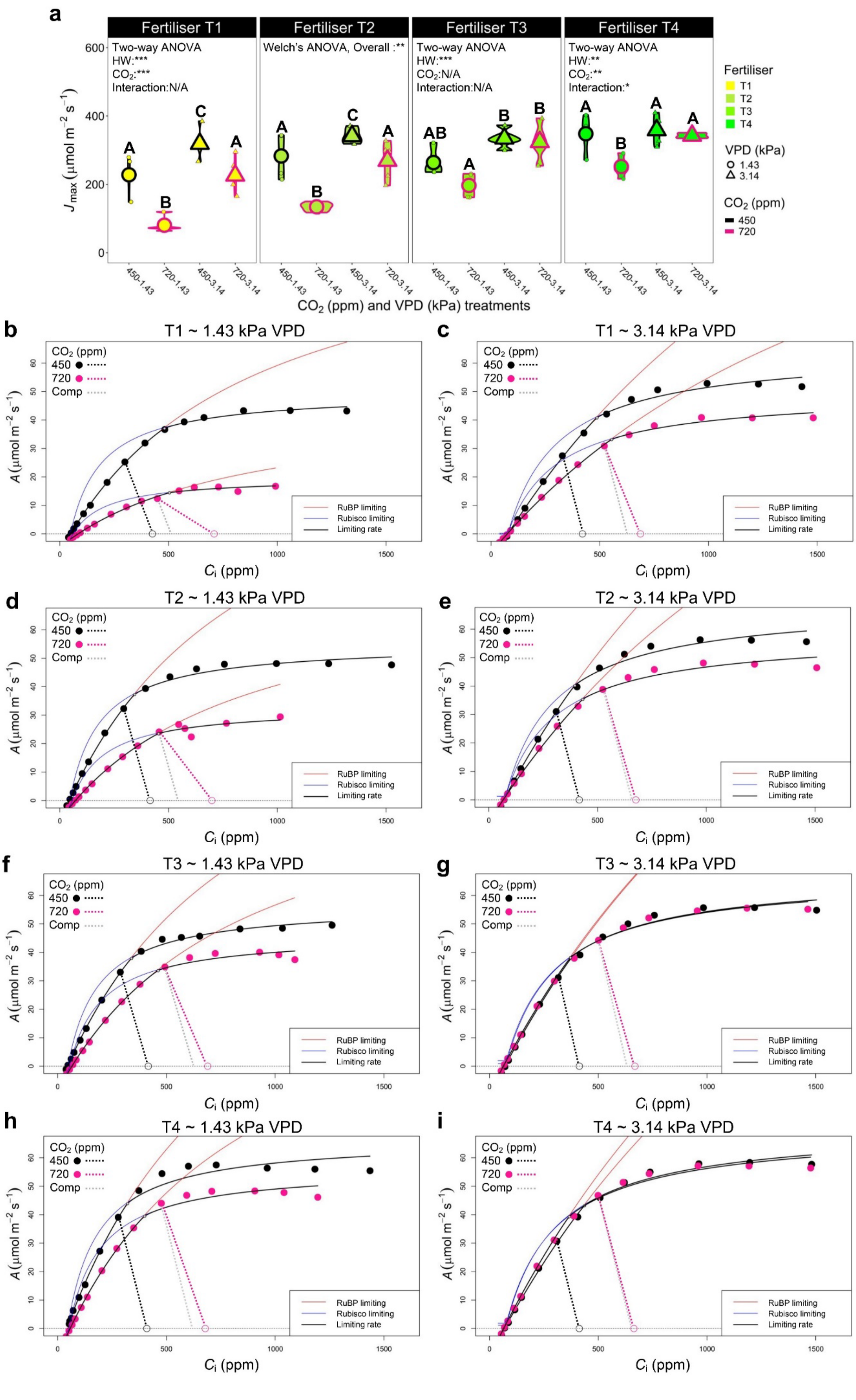
a
Fertiliser T1
Fertiliser T2
Fertiliser T3
Fertiliser T4
Two-way ANOVA
HW:***
CO2:***
Interaction:N/A
Welch's ANOVA, Overall :**
Two-way ANOVA
HW:***
CO2:N/A
Interaction:N/A
Two-way ANOVA
HW:**
CO2:**
Interaction:*
J max (μmol m−2 s−1)
CO2 (ppm) and VPD (kPa) treatments
450-1.43
720-1.43
450-3.14
720-3.14
Fertiliser
T1
T2
T3
T4
VPD (kPa)
1.43
3.14
CO2 (ppm)
450
720
b
T1 ~ 1.43 kPa VPD
c
T1 ~ 3.14 kPa VPD
d
T2 ~ 1.43 kPa VPD
e
T2 ~ 3.14 kPa VPD
f
T3 ~ 1.43 kPa VPD
g
T3 ~ 3.14 kPa VPD
h
T4 ~ 1.43 kPa VPD
i
T4 ~ 3.14 kPa VPD
CO2 (ppm)
450
720
Comp
RuBP limiting
Rubisco limiting
Limiting rate
A (μmol m−2 s−1)
Ci (ppm)

**Extended Data Fig. 5. Further biochemical analyses of wheat plants assessed under different $CO_2$ and vapour pressure deficit (VPD) growth conditions.** (**a**) Maximum rate of photosynthetic electron transport ($J_{max}$) of plants grown and assessed at different $CO_2$ concentrations and VPD scenarios, grouped by fertiliser treatment. (**b-i**) Averaged photosynthesis ($A$)/ intercellular $CO_2$ concentration ($C_i$) curves from the 4 N-fertiliser scenarios across non-heatwave (1.34 kPa) and high-VPD heatwave (3.14 kPa) conditions. Within each graph, supply functions and data points are black for ambient $CO_2$ (450 ppm) treatment pink for high $CO_2$ (720 ppm) treatment. Gray lines represent 450 ppm sample slopes marked immediately adjacent to high $CO_2$ samples as comparisons (comp). The hollow symbols represent $C_a$ at growth room $CO_2$ concentration. $n$ = 4-5. Large symbols in **a** equal sample means. For Two-way ANOVAs, Tukey post-hoc tests were performed. For Welch's ANOVAs, Games-Howell tests were undertaken. Different letters indicate significant differences of $p \leq 0.05$. Asterisks equal, * = $p < 0.05$, ** = $p < 0.01$ and *** = $p < 0.001$.

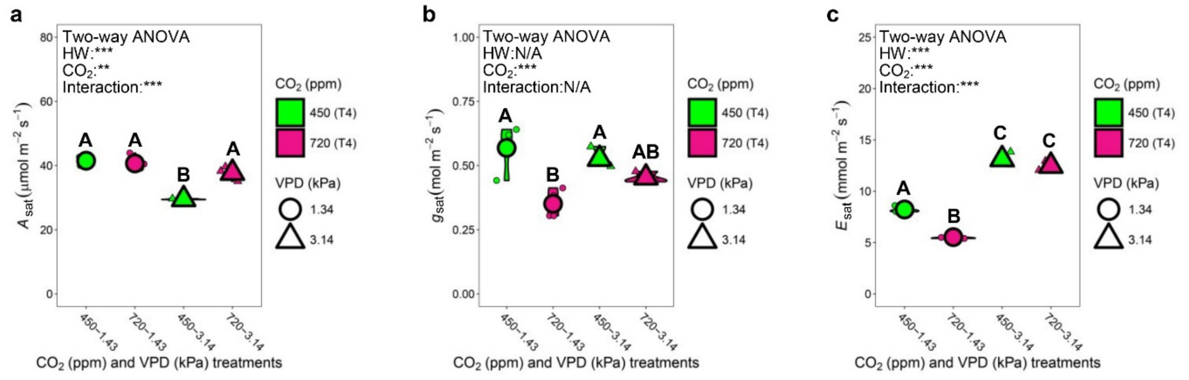


**Extended Data Fig. 6. Plant gaseous exchanges after 5 minutes of light-saturating photosynthesis measurements immediately prior to dark treatment.** Measurements conducted under (**a**-**c**) Saturating light (**a**) photosynthesis ($A_{sat}$), (**b**) stomatal conductance to water vapour ($g_{sat}$) and (**c**) transpiration ($E_{sat}$) using corresponding growth chambers conditions to growth chambers. $n = 3$-4. Large symbols equal means. For Two-way ANOVAs, Tukey post-hoc tests were performed. Different letters within graphs indicate significant differences of $p \leq 0.05$. Asterisks equal, * = $p < 0.05$, ** = $p < 0.01$ and *** = $p < 0.001$.

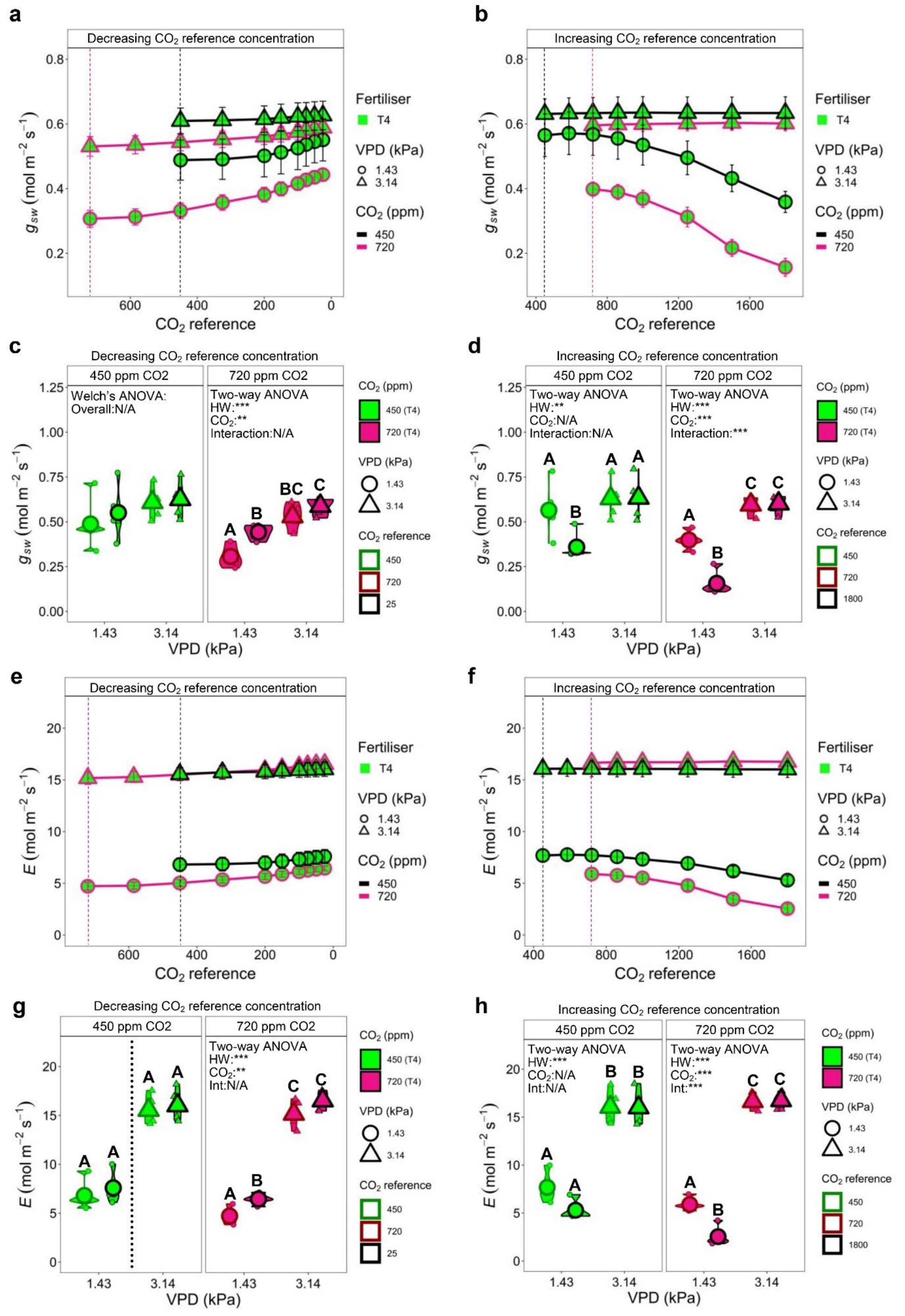


**Extended Data Fig. 7. Plant stomatal conductance to water vapour ($g_{sw}$) and transpiration ($E$) responses to changes in $CO_2$ reference generated from $CO_2$ response curves.** (a-b)

Assessment of $g_{sw}$ changes during $A$/$C_i$ curves from different $CO_2$ and vapour pressure deficit (VPD) growth scenarios. (**a**) Decreasing and (**b**) increasing $CO_2$ reference. (**c**-**d**) Assessment of $g_{sw}$ differences between plants grown at 450 ppm or 720 ppm $CO_2$ concentration. (**c**) Decreasing $CO_2$ reference analysis and (**d**) increasing $CO_2$ reference. Comparisons were between minimum $CO_2$ reference (25 ppm) and growth $CO_2$ reference (450 or 720 ppm), or growth $CO_2$ reference (450 or 720 ppm) and maximum $CO_2$ reference analysis (1800 ppm). (**e**-**f**) Assessment of $E$ changes during $A$/$C_i$ curves from different $CO_2$/ and vapour pressure deficit (VPD) growth scenarios. (**e**) Decreasing and (**f**) increasing $CO_2$ reference. (**g**-**h**) Assessment of $E$ differences between plants grown at 450 ppm or 720 ppm $CO_2$ concentration. (**g**) Decreasing $CO_2$ reference analysis and (**h**) increasing $CO_2$ reference analysis based on same $CO_2$ reference concentrations as in **c**-**d**. $n$ = 4-5. Large symbols equal means. For Two-way ANOVAs, Tukey post-hoc tests were performed. For Welch's ANOVAs, Games-Howell tests were undertaken. Dotted lines in **a**, **b**, **e** and **f** highlight the starting $CO_2$ concentration of the curve. In **g**, the dashed line indicates two one-tailed Student's t-tests were undertaken. Different letters within graphs indicate significant differences of $p \leq 0.05$. Asterisks equal, * = $p < 0.05$, ** = $p < 0.01$ and *** = $p < 0.001$.

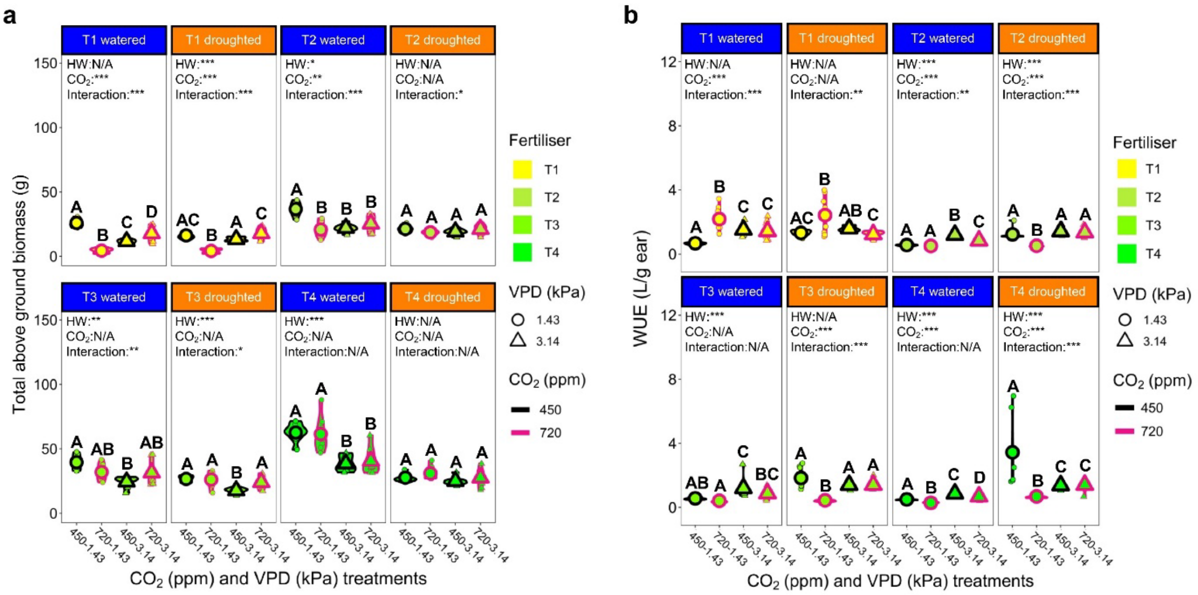


**Extended Data Fig. 8. Total biomass and water-use efficiency (WUE) of wheat plants grown under different $CO_2$, VPD, N-fertiliser regimes with continual irrigation or drought treatment.** (**a**) Total above ground biomass production and (**b**) WUE. WUE calculated by dividing total water application by ear weight. $n = 8$. Large symbols equal means. Statistical testing undertaken using generalised linear models with post-hoc assessment conducted by computing estimated marginal means. Different letters within graphs indicate significant differences of $p \leq 0.05$. Asterisks equal, * = $p < 0.05$, ** = $p < 0.01$ and *** = $p < 0.001$.
